# Kinetic-Mechanistic Evidence for Which *E. coli* RNA Polymerase-λP_R_ Open Promoter Complex Initiates and for Stepwise Disruption of Contacts in Bubble Collapse

**DOI:** 10.1101/2020.09.11.293670

**Authors:** Dylan Plaskon, Kate Henderson, Lindsey Felth, Cristen Molzahn, Claire Evensen, Sarah Dyke, Irina Shkel, M. Thomas Record

## Abstract

In transcription initiation, specific contacts between RNA polymerase (RNAP) and promoter DNA are disrupted as the RNA-DNA hybrid advances into the cleft, resulting in escape of RNAP. From the pattern of large and small rate constants for steps of initiation at λP_R_ promoter at 19°C, we proposed that in-cleft interactions are disrupted in extending 3-mer to 5-mer RNA, −10 interactions are disrupted in extending 6-mer to 9-mer, and −35 interactions are disrupted in extending 10-mer to 11-mer, allowing RNAP to escape. Here we test this mechanism and determine enthalpic and entropic activation barriers of all steps from kinetic measurements at 25°C and 37°C. Initiation at 37°C differs significantly from expectations based on lower-temperature results. At low concentration of the second iNTP (UTP), synthesis of full-length RNA at 37°C is slower than at 25°C and no transient short RNA intermediates are observed, indicating a UTP-dependent bottleneck step early in the 37°C mechanism. Analysis reveals that the 37°C λP_R_ OC (RP_O_) cannot initiate and must change conformation to a less-stable initiation complex (IC) capable of binding the iNTP. We find that IC is the primary λP_R_ OC species below 25°C, and therefore conclude that IC must be the I_3_ intermediate in RP_O_ formation. Surprisingly, Arrhenius activation energy barriers to five steps where RNAP-promoter in-cleft and −10 contacts are disrupted are much smaller than for other steps, including a negative barrier for the last of these steps. We interpret these striking effects as enthalpically-favorable, entropically-unfavorable, stepwise bubble collapse accompanying disruption of RNAP contacts.

**Significance:** Transcription initiation is highly regulated. To understand regulation, mechanisms of initiation and escape of RNA polymerase (RNAP) from the promoter must be understood. RNAP forms a highly-stable open complex (RP_O_) with λP_R_ promoter at 37°C. From experiments determining effects of temperature on rate constants for each step of RNA synthesis, we find that RP_O_ cannot bind the initiating nucleotides, that the I_3_ intermediate and not RP_O_ is the initiation complex, and that contacts of RNAP with single-stranded DNA of the discriminator and −10 region and with −35 duplex DNA are disrupted stepwise as the RNA-DNA hybrid moves into the cleft. Evidence is obtained for stepwise bubble collapse and base stacking accompanying disruption of interactions of the single-stranded discriminator and −10 regions with RNAP.

## Introduction

Transcription initiation, including open complex formation and subsequent steps of NTP binding and synthesis of a RNA-DNA hybrid, is a prime target for regulation. An understanding of the kinetics and mechanism of transcription initiation (1–9) is needed to complement structural characterization of RNAP-promoter initiation complexes (10–13), with applications to antibiotic development and synthetic promoter design (14). In the initial specific closed complex (RP_C_), the start site region of duplex promoter DNA is not contacted by RNAP (10, 15). Recent kinetic-mechanistic and structural research have increased our understanding of the mechanisms by which the start site region is opened by RNAP using binding free energy and the template strand is brought to the vicinity of the RNAP active site to form the initial (unstable) open complex (I_2_) (11, 15–22).

At numerous promoters, including λP_R_, much more stable open complexes are formed from I_2_ by conformational changes that appear to be directed by interactions of σ^70^ region 1.2 with the non-template (nt) discriminator strand in the cleft (23). At λP_R_, these interactions directs downstream mobile elements (DME) to assemble and tighten on the downstream duplex after opening the initiation bubble and rotating the downstream duplex by about one turn (360°) to form I_2_. At 37 °C these DME interactions convert unstable I_2_ to the very stable λP_R_ RP_O_ complex via the I_3_ intermediate (1, 15, 24–27), a process which is strongly favored by increasing temperature (13 °C - 42 °C). MnO_4_-footprinting revealed that the same bubble region was open in λP_R_ I_2_ as in the stable λP_R_ OC at 10 °C, and that the reactivity of the +1 thymine on the template strand was similar in these OC, indicating that the positioning of the +1 thymine might be the same (28). The question of which OC is/are capable of initiation upon NTP addition has not been addressed previously.

In the steps of productive initiation, translocation of the RNA-DNA hybrid into the cleft disrupts specific contacts of RNAP with in-cleft, −10, and −35 regions of the promoter, resulting in collapse of (and duplex formation by) the upstream portion of the initiation bubble (−1 to −11 for λP_R_), escape of RNAP from the promoter, and dissociation of the σ^70^ subunit (2, 29, 30). Escape of RNAP from the λP_R_ occurs after synthesis of an 11-mer RNA (5, 31).

At the λP_R_ promoter, stress buildup and disruption of RNAP-promoter contacts have been monitored by their effects on the composite second order rate constant k_i_ (k_cat_/K_M_ analog) for the individual steps of NTP addition up to promoter escape (2 ≤ i ≤ 11) at 19 °C. Values of k_i_ includes contributions from reversible translocation, reversible NTP binding and irreversible catalysis (32). Values of k_i_ for steps of initiation involving translocation divide into three groups. Values of k_I_ are much larger for synthesis of 3-mer, 6-mer, and 10-mer RNA than for the other six steps (32).

Elongation kinetic studies reveal that the translocation step before NTP binding is rapidly reversible and that the post-translocated state is intrinsically less stable than the pre-translocated state (33–35). The post-translocated state is stabilized by binding of the next NTP (32, 35). In initiation, effects of translocation stress greatly increase the bias toward the pretranslocated state (32). Large k_i_ values indicate elongation steps where translocation is less unfavorable because stress has not yet built up (2-mer → 3-mer) or where stress has been released by disruption of contacts in previous steps (5-mer → 6-mer; 9-mer → 10-mer). We proposed that moderate-strength in-cleft interactions are disrupted primarily in steps 4-5, strong −10 interactions are disrupted primarily in steps 7-9, and relatively weak −35 and upstream interactions are disrupted in step 11, resulting in escape of RNAP (32).

Here we report the kinetics of initial transcription at the λP_R_ promoter at 25 °C and 37 °C. Rapid-quench mixing is used to determine overall rates of full-length RNA synthesis and rate constants for individual nucleotide addition steps at two nucleotide conditions (designated high UTP, low UTP) for comparison with the 19 °C results. These studies address the questions of which OC conformation is capable of binding nucleotides and which specific RNAP-promoter contacts are disrupted in the individual steps of hybrid-translocation, resulting in collapse of the initiation bubble and escape of RNAP from the promoter. In addition, they provide key thermodynamic information about the I_3_ OC and the steps by which the unstable I_2_ OC converts to RP_O_.

## Results

### Unexpected Differences Between ÁP_R_ Promoter Initiation Kinetics at 37 °C and 25 °C

Time courses (≤ 0.5 s to ≥ 90 s) of transcription initiation by *E. coli* RNA polymerase (RNAP) at the λP_R_ promoter at 37 °C and 25 °C were obtained by rapid mixing at two different sets of NTP concentrations for comparison with previous results at 19 °C (32). The competitor heparin, added with the NTP mixture, ensures that FL RNA synthesis is single-round by preventing re-initiation by any dissociated RNAP. The panels of Fig. 1 show polyacrylamide gel electrophoresis (PAGE) separations of RNAs present in samples quenched at a series of time points during initiation and the transition to elongation. Amounts of each RNA length present at each time are detected by incorporation of ^32^P-GTP or ^32^P-UTP. Fig. 1 panels A-C are for a “high UTP” condition (final concentrations 200 μM UTP and ATP, 10 μM GTP, 17 nM ^32^P-GTP), and panels D-F are for a “low UTP” condition (final concentrations 10 μM UTP, 200 μM ATP and GTP, 17 nM ^32^P-UTP). Efficient incorporation of α-^32^P label into the transcript is achieved by use of a low concentration (10 μM) of the corresponding unlabeled NTP. The key first step of initiation (pppApU synthesis) is of course favored by the high UTP condition as compared to the low UTP condition.

**Figure 1:**
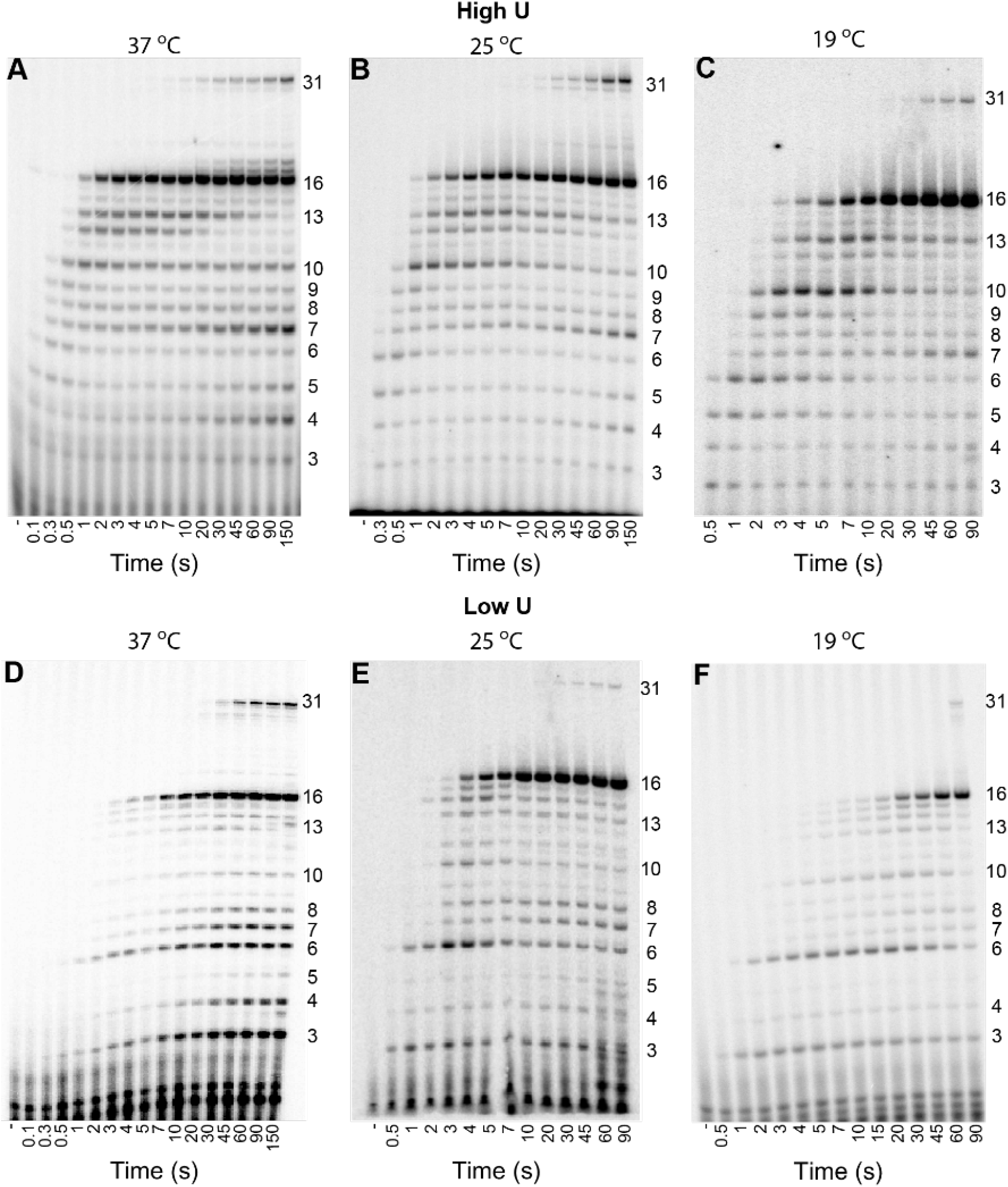
High-Resolution Time Courses of Transcription Initiation from λP_R_ Promoter as a Function of Temperature. Time-dependence of amounts of individual short and long RNA products of transcription initiation at the λP_R_ promoter at 37 °C (**Panels A, D**) and 25 °C (**Panels B, E**), compared with 19 °C (**Panels C, F**; from ref (32)). Polyacrylamide gel separations are shown for the indicated times (0.1s to 150s) after adding NTPs and heparin to premixed RNAP and λP_R_ promoter DNA. In all cases CTP is omitted to stop transcription from this modified λP_R_ ITR at a 16-mer. **Panels A-C**: 200 μM ATP and UTP, with 10 μM GTP. RNA is labelled by addition of ~17nM α-^32^P-GTP. **Panels D-F**: 200 μM ATP and GTP, with 10 μM UTP. The RNA is labelled by addition of ~17nM α-^32^P-UTP. Slow formation of longer transcripts is likely the result of misincorporation (32).

For these experiments, the initial transcribed sequence of wild-type λP_R_ was modified to pppA^+1^pUpGpUpApG^+6^pUpApApGpG^+11^pApGpGpUpU^+16^pC so the first C in the transcript occurs at position +17 and transcription pauses at a 16-mer when CTP is withheld. This pause occurs after escape of RNAP, which for this promoter is deduced to occur at the 10-mer to 11-mer step (5). All RNAs greater than 10-mer in length are considered part of the full-length (FL) RNA population. The transient accumulation of 12-mer and 13-mer may result from the reduction in rate constants for the subsequent steps caused by coupling of translocation to disruption of σ^70^ - core RNAP contacts. The next C in the transcript is near the end of the fragment at +32, so read-through at +17 leads to a second pause at a 31-mer. Transcription occurs slowly near the end of a fragment, so this second pause is effectively a stop point.

Effects of temperature on three different aspects of initiation are shown in these gels for the two NTP conditions examined: 1) transient buildup and decay of short (3-mer to 10-mer) RNA intermediates in FL RNA synthesis by productive OC, observed at short times (<20 s) after NTP addition; 2) buildup of FL RNA (>10-mer) after an initial lag; and 3) synthesis at short times (<20 s) of an initial short RNA by non-productive OC, often but not always followed by slower release of that short RNA and re-initiation (abortive synthesis).

At an overview level, Fig. 1 shows that patterns of RNA synthesis by both productive and nonproductive complexes are similar at 25 °C and 19 °C, and that rates of steps of productive initiation and FL RNA synthesis as well as abortive initiation by nonproductive complexes all are larger at 25 °C. However, Fig. 1 reveals some large differences at 37 °C in rate and/or pattern of RNA synthesis by productive and nonproductive complexes from expectation based on the lower temperature results.

### Full-Length (FL) RNA Synthesis

Fig. 1 reveals that, while the time required for synthesis of FL (>10-mer) RNA at the high UTP condition (panels D-F) decreases monotonically with increasing temperature, the time required for FL synthesis at 37 °C at the low UTP condition (panels A-C) is clearly greater than at 25 °C. Normalized amounts of FL RNA per OC are plotted vs time (log scale) in Fig. 2, which compares FL RNA synthesis at 25 °C and 37 °C with previous results at 19 °C for high UTP (Panel A) and low UTP (Panel B) conditions. Results are averages of 2-4 independent experiments including those in Fig. 1. Representative sets of unnormalized data from single experiments are plotted in Fig. S1. Plateau amounts of FL RNA per OC are 0.5 ± 0.15 RNA/OC (e.g. Fig. S1), demonstrating that ~50% of the population of open complexes make a FL RNA, and are termed productive complexes. Each experiment is normalized relative to its plateau value before averaging.

**Figure 2:**
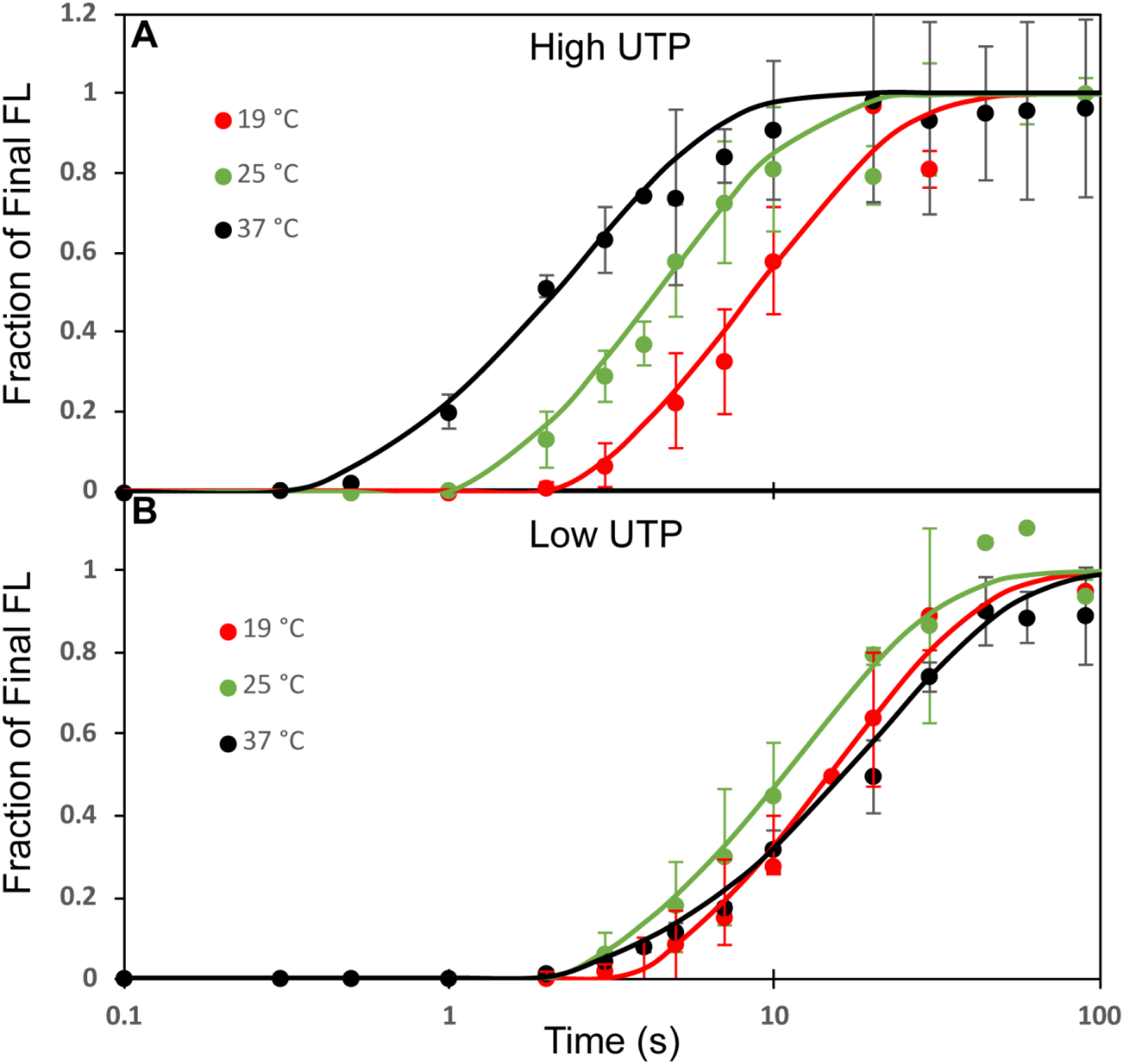
Kinetics of FL RNA Synthesis in Single-Round Initiation as a Function of Temperature. Average total amount of FL (length >10 bp) RNA synthesized as a function of time at 19 °C (red), 25 °C (green), and 37 °C (black) from 2-4 experiments like those in Figure 1 at 10 μM GTP (**Panel A**) and 10 μM UTP (**Panel B**). Amount of FL RNA is normalized to the amount of FL produced by 150 s for each condition. Error bars represent one standard deviation.

At each condition, after an initial lag, FL RNA synthesis occurs over a ~20-fold time interval, from ~2 s to ~40 s at the low UTP condition at all temperatures and also at the high UTP condition at 19 °C, from ~1 s to ~20 s at the high UTP condition at 25 °C, and from ~0.4 s to ~8 s at the high UTP condition at 37 °C. For high UTP conditions (Fig. 2A), as the temperature is increased the length of the lag phase decreases and the rate of subsequent FL RNA synthesis increases. For low UTP (high GTP) conditions (Fig. 2B) the lag is longer and less temperature-dependent and the rate of FL synthesis is slower than at high UTP (Fig. 2A; low GTP), even though the RNA at the point of escape has more G bases (4) than U bases (3). Most notably, while FL synthesis is faster at 25 °C than at 19 °C at low UTP (Fig 2B) as well as at high UTP (Fig 2A), FL synthesis at 37 °C and low UTP is as slow or slower than at 19 °C.

As observed previously (5), the kinetics of FL synthesis after the initial lag are well-described as first-order (single exponential) approaches to the plateau value characterized by a first-order rate constant k_FL_. At high UTP, k_FL_ increases monotonically with temperature, with a positive Arrhenius activation energy that decreases with increasing temperature (Fig. S2). At low UTP, all values of k_FL_ are smaller and the 25 °C value is larger than the 37 °C value, corresponding to a negative activation energy between 25 °C and 37 °C (Fig. S2) and indicating a change in the rate determining step(s) with increasing temperature.

### Transient Short RNA Intermediates in FL RNA Synthesis; Separation from Abortive RNA

At 25 °C, prior to and during the appearance of FL RNA (< 20 s after NTP addition), transient buildups and decays in amount of various short on-pathway RNAs of increasing length (3-mer to 10-mer) are observed in the gels of Fig. 1, as previously reported at 19 °C (5). The time evolution of amounts of four different short RNAs (3-mer, 5-mer, 6-mer, 10-mer) in initiation at high and low UTP conditions at 37 °C and 25 °C are compared with previous results for 19 °C in Fig. 3. Results plotted are averages obtained from analysis of multiple gels like in Fig. 1 and are normalized per open complex before averaging.

**Figure 3:**
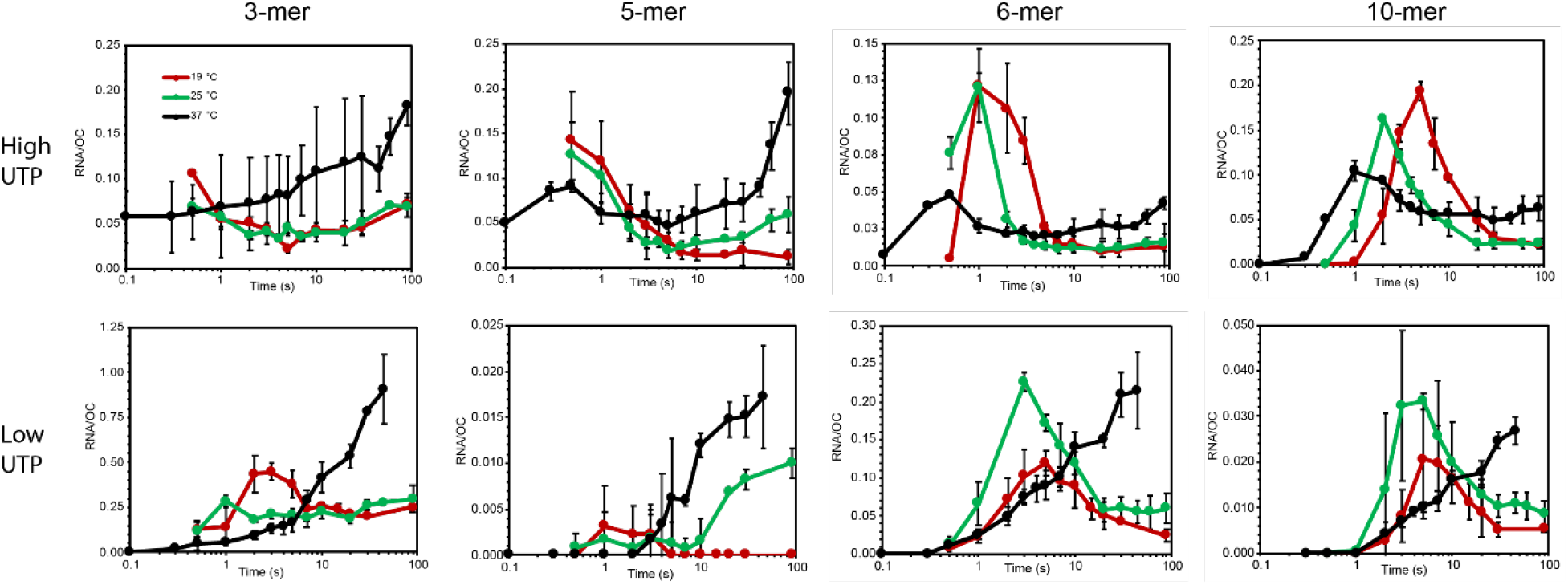
Time Courses of Short RNA Populations in Initiation as a Function of Temperature. **Panels A** (10 μM GTP) and **B** (10 μM UTP) compare time courses (log scale) of synthesis of transient RNAs by productively-initiating complexes and of non-productive complexes at 19 °C, 25 °C, and 37 °C. Amounts of short RNA present are normalized per open complex (OC) and plotted vs time. Each point is an average of 2-4 experiments.

Amounts of each RNA plotted vs. time in Fig. 3 include both transient RNA intermediates on the pathway to FL RNA synthesis by productive OC and short RNAs synthesized by nonproductive complexes (32). To emphasize the short time range where FL RNA is being synthesized by productively-initiating complexes, while also showing the longer time behavior, these plots use logarithmic time axes. Linear time scale plots of these data and data for other short RNAs (4-mer to 9-mer) observed at each temperature and NTP condition, which display more clearly the slow kinetics of abortive initiation by nonproductive complexes and the separation of intermediates in productive initiation from abortive products, are shown and discussed briefly in Supplemental as Figs. S3-S6 and accompanying text. Extrapolation of the amount of abortive RNA back to short times (SI Figs. S3-S6) provides a baseline to quantify the amount of each short-RNA transient as a function of time in FL RNA synthesis by productive complexes.

At 25°C, at both low UTP and high UTP, Fig. 3 shows that amounts of all four RNA species increase rapidly and then decrease in the first 10 s, consistent with previous observations at 19 °C and indicating significant transients on the pathway to productive synthesis (32). Each transient occurs earlier at 25 °C than at 19 °C at both NTP conditions, as expected since most reaction rates increase with increasing temperature. Also each transient occurs earlier at high UTP than at low UTP at both temperatures, expected because UTP is a reactant in the first step of initiation (pppApU synthesis).

At 37 °C, however, no significant transient population of any intermediate is observed in FL RNA synthesis at low UTP. At high UTP, 37 °C, transient populations of some longer RNA intermediates (5-mer, 6-mer, 9-mer, 10-mer) are detected at times similar to those observed at 25 °C and high UTP, but the amounts are small by comparison to what is observed at lower temperatures. These observations, and the slower synthesis of FL RNA at low UTP at 37 °C than at 25 °C, all indicate that the population of 37 °C LPR OC, unlike OC populations at lower temperatures, is unable to bind the initiating NTP and synthesize pppApU initiate without undergoing a conformational change. This mechanistic step is introduced in the analysis of the initiation kinetics below.

The stability of the λP_R_ OC is more than 30-fold greater at 37 °C than at 19 °C (22, 25). We previously proposed that the population distribution of λP_R_ OC also changed with temperature, shifting from the very-stable 37 °C RP_O_ with strong downstream interactions between RNAP DME and duplex DNA extending to +20 to a mixed population of RP_O_ and the less stable intermediate OC designated I_3_ at lower temperature (25). If only I_3_ and not RP_O_ can bind the two initiating NTP, then the differences in rates of FL RNA synthesis between 37 °C and lower temperatures are readily explained. In Analysis and Discussion this proposal is incorporated into the initiation mechanism and used to analyze the kinetics of transient and FL RNA synthesis by productive complexes.

Figs. 1, 2 and S3-S6 also show the complex effects of temperature and NTP condition on initial and multi-round (abortive) synthesis of short RNAs by nonproductive complexes that stall before the escape point. These are discussed briefly in Supplemental.

## Analysis and Discussion

### Evidence for an OC Conformational Change Prior to NTP Binding at 37 °C but not at 19 °C

Previously, 19 °C λP_R_ initiation kinetic data were fit to Mechanism 1 (Fig. 4) which begins with ordered, reversible binding of the substrates (ATP (+1), UTP (+2)) to the OC, followed by irreversible catalysis to synthesize the dinucleotide pppApU. In this fitting, individual values of the UTP binding constant (1/K_m_ analog) and the catalytic rate constant (k_cat_ analog) were obtained because the UTP binding constant in pppApU synthesis is relatively large (3 x 10^4^ M^−1^), leading to near-saturation of ATP-bound OC with UTP at 200 μM UTP. No evidence was obtained for a conformational change in the OC prior to NTP binding (32).

**Figure 4:**
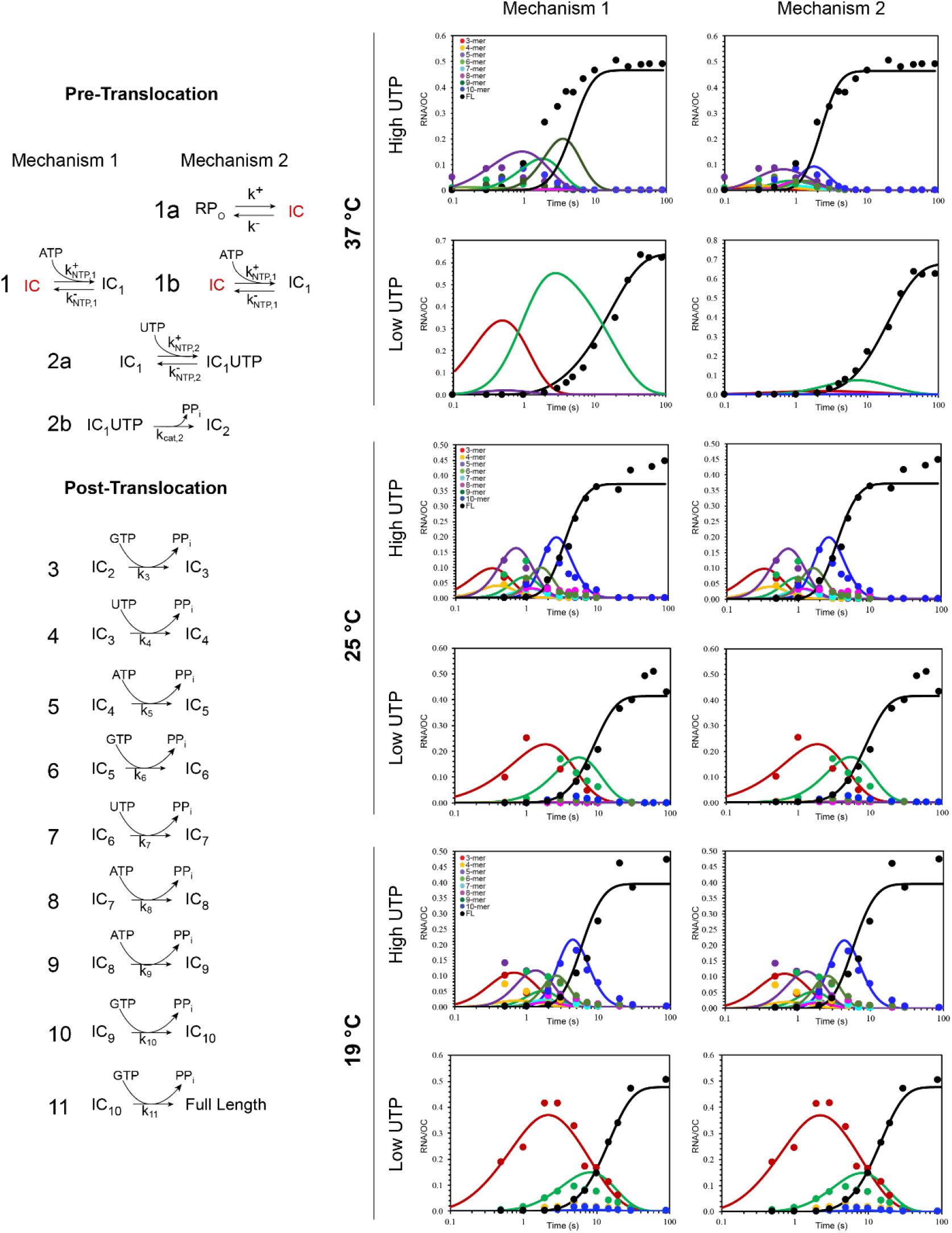
Comparison of Fits of Initiation Kinetic Data to Two Mechanisms. **LEFT:** Comparison of two mechanisms used to fit the λP_R_ transcription initiation data. In **Mechanism 1,** the first nucleotides (A, U) bind to the preformed active site without any conformational changes in the OC. This mechanism provides an excellent fit to 19 °C initiation kinetic data (32). **Mechanism 2:** To fit λP_R_ initiation data at 37 °C, an initial, reversible, highly temperature dependent OC rearrangement (step 1a) must be added to **Mechanism 1** in which the stable λP_R_ OC (RP_O_) is converted to a less stable but initiation competent OC form (IC, red). Formation of IC from RP_O_ at 37 °C is followed by reversible ATP binding at the transcription start site (step 1b), reversible UTP binding at position +2 (step 2a) and irreversible catalysis to form the initiating dinucleotide (step 2b). Subsequent steps of both mechanisms are the same, each including reversible translocation, reversible NTP binding, and irreversible catalysis. They are described at the NTP concentrations investigated here by a composite second order rate constant k_i_ (k_cat_/K_m_ analog; (32)). **RIGHT**: Simulations of time evolution of the population of transient intermediates and full-length RNA synthesis by productive complexes, compared with experimental data. Panels show results for 37 °C (top), 25 °C (middle), and 19 °C (bottom) and two nucleotide conditions (high UTP (top), low UTP (bottom)). Fits to Mechanism 1 are shown in the left column and clearly do not describe the experimental data at 37 °C. Fits to Mechanism 2 are much more accurate at 37 °C, and achieve nearly identical quality of fit at 19 °C and 25 °C, providing values of the equilibrium constant for the conformational change RP_O_ → IC at each temperature.

Every subsequent step of initiation of RNA synthesis begins with reversible translocation. Translocation in elongation is intrinsically unfavorable (32, 35, 36), presumably because of disruption of one RNA-DNA hybrid base pair (a downstream DNA-DNA base pair is disrupted but an upstream DNA-DNA base pair is formed). Translocation in initiation is more unfavorable (32). In addition to disruption of one DNA-DNA base pair, translocation stress accumulates and RNAP-promoter contacts are disrupted in many of these steps. Because translocation equilibrium constants for initiation steps are small, the overall equilibrium constant for the reversible translocation and NTP binding steps (analog of 1/K_m_) cannot be separated from the rate constant of the irreversible catalytic step (k_cat_) for the NTP concentrations investigated (32). Hence each step of RNA-DNA hybrid extension up to the predicted point of escape is accurately quantified using a composite second order rate constant k_i_ (the analog of k_cat_/K_m_; see (32)). This simplification of the enzyme mechanism is shown as Mechanism 1 in Fig. 4, together with the fit of the transient RNA peaks and FL RNA synthesis at 19 °C determined previously for low and high UTP conditions.

Also shown in Fig. 4 are the results of fitting the kinetic data at 25 °C and 37 °C to Mechanism 1 at low and high UTP conditions. At 25°C, rate constants k_i_ obtained from these fits accurately reproduce the transient appearance and disappearance of on-pathway short RNAs and the kinetics of synthesis of FL RNA. However, it is clear from Fig. 4 that Mechanism 1 is not sufficient to describe the kinetics of productive initiation at 37 °C. To obtain an accurate fit to the 37 °C kinetic data requires the addition of an unfavorable reversible step at the beginning of the mechanism, prior to initial ATP binding (Fig. 4, Mechanism 2, step 1a). This step is a conformational change in the OC, which we propose is the conversion of the very stable OC (RP_O_) to another OC conformation that we designate the initiation complex (IC), characterized by the equilibrium constant K_1a_ (for RP_O_ → IC). All other steps of Mechanism 2 are the same as Mechanism 1. Fig. 4 also shows that use of Mechanism 2 to fit 19 °C and 25 °C data sets does not affect the quality of these fits and yields estimates of K_1a_ at these temperatures.

These fits predict that equilibrium constant K_1a_ is extremely temperature dependent, greatly favoring RP_O_ at 37 °C (K_1a_ ≈ 0.01; 99% RP) but favoring IC at 19 °C (K_1a_ ≈ 5.3; more than 80% IC). A near-equimolar ratio of RP_O_ and IC is predicted at 25 °C (K_1a_ ≈ 0.70; ~40% IC; ~60% RP_O_). Good fits of 19 °C and 25 °C kinetic data to Mechanism 1 are obtained because a significant fraction of the OC population is initially in the IC conformation at these temperatures. The strong decrease in K_1a_ with increasing temperature indicates that the enthalpy change for the conversion of RP_O_ to IC is large in magnitude and negative; van’t Hoff analysis yields 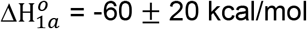. The standard free energy change 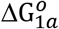 for this conversion ranges from ~ 2.7 kcal at 37 °C to −1 kcal at 19 °C, and the corresponding entropy change 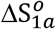 is −200 ± 60 eu. Conversion of RP_O_ to IC therefore shows near-complete enthalpy-entropy compensation, like many other protein processes.

### Only the I_3_ Intermediate OC and not RP_O_ Initiates Transcription from λP_R_ Promoter Upon NTP Addition

Evidence exists for two open intermediates (I_2_, I_3_) on the pathway to formation of the RP_O_ complex at the λP_R_ promoter. The thermodynamic, kinetic and footprinting information available for these intermediates support the proposal that IC is I_3_. I_2_, the OC formed in the DNA opening step, is unstable at all temperatures (25). It is unstable with respect to I_3_ and/or RP_O_ at higher temperatures and unstable with respect to closed complexes at lower temperatures. Conversion of I_2_ to I_3_ involves folding of 100-150 amino acid residues of RNAP DME (25), and is thought to strengthen contacts between the proximal downstream duplex and the β lobe and the β’ clamp (27). Conversion of I_3_ to RP_O_ is thought to involve primarily an interaction of the downstream jaw and associated DME with the distal downstream duplex (+10 to +20), which serves to tighten the entire RNAP-promoter interface in the OC. The OC formed by the jaw deletion variant of RNAP and by downstream truncation variants of the promoter are thought to be models of I_3_; equilibrium constants for forming these variant OC are one to two orders of magnitude smaller than binding of WT RNAP to full-length promoter DNA at 37 °C. Hydroxyl radical footprinting of the open complex with the jaw deletion RNAP variant reveals that the entire RNAP-promoter interface in the OC is less protected and hence “looser” and more hydrated than in the WT RNAP OC (27).

From this body of previous research, the stable OC population was proposed to be an equilibrium mixture of I_3_ and RP_O_, with RP_O_ highly favored at 37 C and I_3_ increasing in significance at lower temperature (15, 27), but the details of this were not known. Here we find that the stable OC population is an equilibrium mixture of RP_O_ and the IC initiation complex, with RP_O_ favored at 37 C and IC favored below 25 °C, indicating that IC is I_3_. In support of this, extrapolation of K_1a_ (Mechanism 2) to lower temperature assuming a temperature-independent enthalpy change for RP_O_ → IC 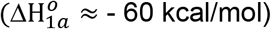 predicts that the IC:RP_O_ population distribution for λP_R_ at 10 °C, before NTP addition, is ~99% IC and only 1% RP_O_. At 10 °C, Gries *et. al.* (28) determined MnO_4_- footprints of both strands of the open region in the stable λP_R_ OC, now identified as the IC initiation complex. In addition, salt-upshifts were used to rapidly destabilize the 10 °C λP_R_ OC and obtain a burst of I_2_, the least stable open intermediate, for MnO_4_- footprinting. Hence the 10 °C λP_R_ OC population, identified in the current research as 99% IC, is more stable and hence more advanced than I_2_ at 10 °C and therefore must be I_3_.

The overall enthalpy change 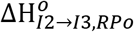, for conversion of I_2_ to the I_3_ – RP_O_ equilibrium mixture at temperatures from 7 °C to 37 °C can be estimated by comparison of activation energies for OC dissociation (37) and for the DNA closing step (25). 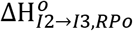 decreases from ~ 40 kcal at 10 C to ~ 7 kcal at 37 C. Identifying IC as I_3_ provides the enthalpy change 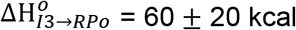, with no detectable temperature dependence. Using 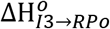 to interpret 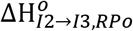, yields values of 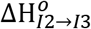 which are very strongly temperature dependent, decreasing from approximately 40 k_cal_ near 10 °C to −50 kcal near 37 °C. Hence the heat capacity change 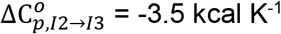. Interpreted in terms of coupled folding, 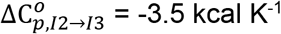 corresponds to folding of 100-150 amino acid residues in the conversion of I_2_ to I_3_. Over 100 conserved residues in this region of the C terminus of β’ are predicted to be intrinsically disordered in free RNAP by the computer algorithm PONDR (26). This prediction from 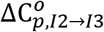 is consistent with the amount of folding predicted from the urea dependence of the dissociation rate constant kd (25). The urea dependence of kd was found to be the same at 10 °C and 37 °C, indicating that all the folding in these steps is in the conversion of I_2_ to I_3_, consistent with the absence of a detectable heat capacity change for the conversion of I_3_ to RP_O_.

### Large, Systematic Changes in Rate Constants, Arrhenius Activation Energies and Transition State Barrier Heights 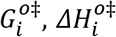, 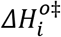, and 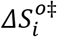 for Steps of Initiation

Composite rate constants k_i_ for the translocation-dependent steps in initial transcription show the same pattern at 25 °C and 37 °C as previously observed at 19 °C, with three distinct regions of small k_i_ that are separated by single steps with larger k_i_ (Figure 5; Table S2). To transition from initiation to elongation, the specific contacts between RNAP and promoter DNA that are essential to form the CC ensemble and direct opening of the transcription bubble must be broken. Previously we interpreted the pattern of 19 °C rate constants k_i_ (k_cat_/K_m_ analog containing the equilibrium constants for translocation and NTP binding and the catalytic step k_cat_) in terms of the serial disruption of these promoter contacts. In this interpretation, differences in k_i_ values arise from differences in the equilibrium constant for translocation at each nucleotide addition step, K_i,trans_. The very similar patterns of low and high k_i_ values at all three temperatures investigated indicate that interactions of RNAP with discriminator, −10 and −35 regions of the promoter are broken in the same steps at all three temperatures.

**Figure 5:**
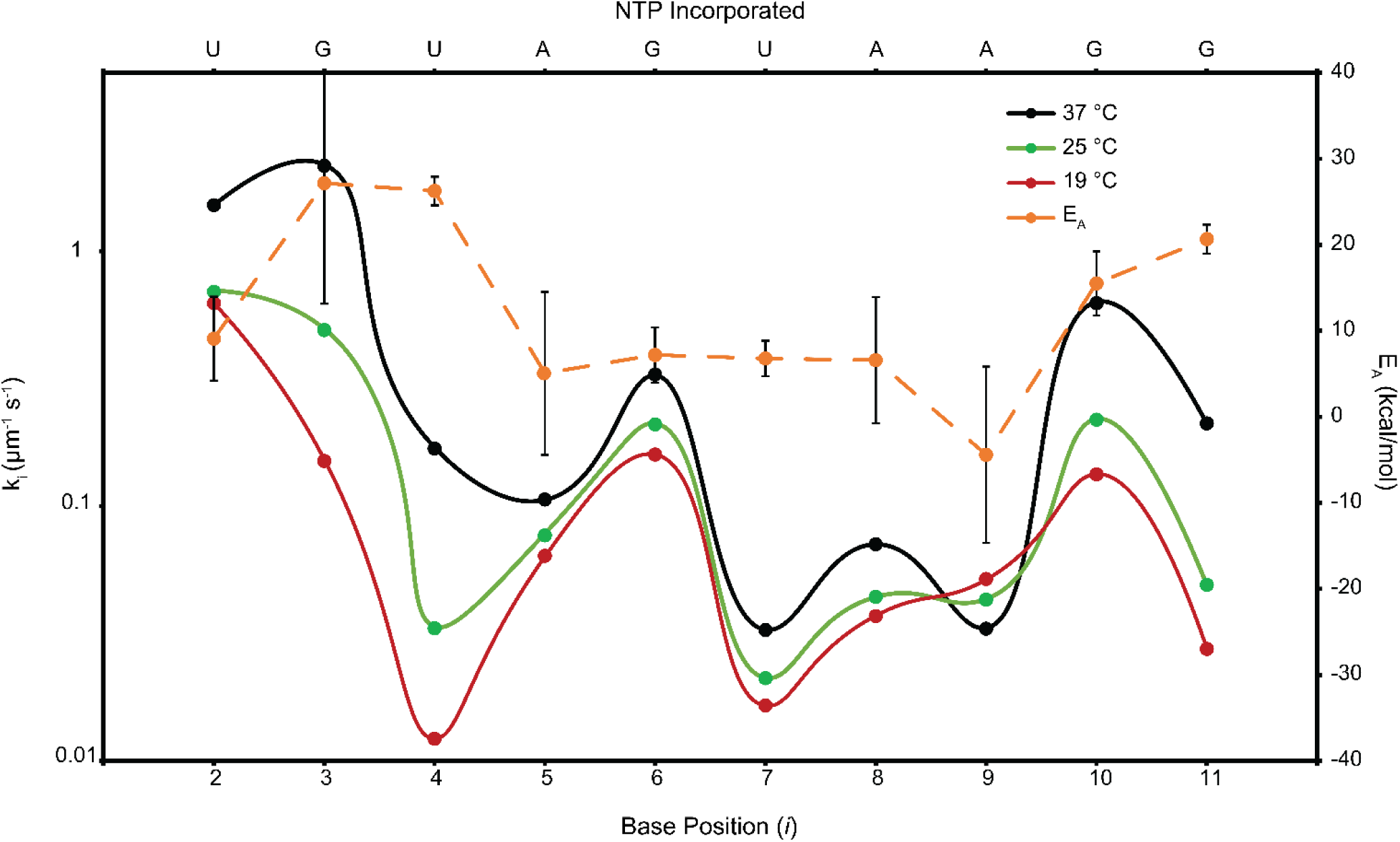
Rate Constants and Activation Energies on the Pathway to Promoter Escape. The composite rate constant (k_i_, in μM^−1^ s^−1^) including reversible translocation (steps 3-11), reversible NTP binding (steps 2-11), and irreversible catalysis (steps 2-11) for each RNA extension step up to the predicted point of escape are plotted vs the extension step for 19 °C (red), 25 °C (green), and 37 °C (black). The pattern of large k_i_ (steps 3, 6, 10) and small k_i_ (steps 4-5, 7-9, and 11) are consistent for each temperature. Arrhenius activation energies (E_A,I_; orange, right scale) of the steps of initiation, determined from the temperature dependence of of k_i_ (Fig. S7), are also shown.

Analysis of the temperature dependences of the k_i_ values yields Arrhenius activation energies (E_A,i_, Fig. 5; Table S3) for the individual steps of initiation. Strikingly, values of E_A,I_ vary by more than 30 kcal (from +27 kcal to −4 k_cal_). E_A,2_ for incorporation of UTP into pppApU, not involving translocation stress, is ~12 kcal, similar to that reported previously for elongation by *E. coli* RNAP (10 – 13 kcal; (36)). E_A,3_ and E_A,4_ for synthesis of 3-mer and 4-mer RNAs are much larger (~26 kcal), while E_A_ values for the following six steps are smaller (~ 6 kcal for synthesis of 5-mer to 8-mer RNA, ~ −4 kcal for 9-mer, and ~ 16 kcal for 10-mer). From Figure 5, k9 decreases with increasing temperature, resulting in a negative E_A,9_. Four of these six steps were previously proposed as steps in which in-cleft and −10 region contacts are disrupted. E_A,11_ is larger than the previous six E_A_ values, though not as large as E_A,3_ and E_A,4._

Significantly, steps with unusually small E_A,I_ values (Fig. 5) follow steps with small rate constants, in which RNAP-contacts with the discriminator and −10 region are disrupted, with an offset of one step. These observations can be explained as the result of stepwise base stacking (and/or base pairing) after stepwise disruption of RNAP contacts with the strands of the discriminator and −10 region in productively initiating complexes, as discussed below.

From rate constants k_i_ and activation energies E_A,I_, the quasi-thermodynamic quantities 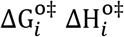, and 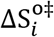 for conversion of reactants to the catalytic transition state of each step can be estimated if the hypothetical maximum rate constant k_max_ for this process (at 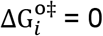) and its temperature dependence are known. As summarized in SI, for this overall 2^nd^ order process we assume the same orientation-corrected diffusion-limited k_max_ ≈ 10^3^ μM^−1^s^−1^ for each step, and an activation energy EA,diff for a diffusion-limited step of 5 kcal mol^−1^ (38). Although the uncertainty in k_max_ is probably one order of magnitude, this is of no consequence for the analysis in Tables S3-S4 and below as long as k_max_ has the same value for each step.

For k_max_ ≈ 10^3^ μM^−1^s^−1^, 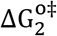 for incorporation of the initiating UTP into pppApU is 4.3 kcal, and values of 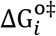 for subsequent steps of initiation (3 ≤ I ≤ 11) are in the range 5.1 kcal to 6.6 kcal (Table S3). Values of 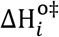 and 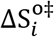 vary over much wider ranges. Like E_A,i_, values of 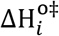 span a range of 30 kcal while values of 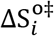 span a range of more than 100 eu (Table S3). For incorporation of the initiating UTP into pppApU, 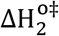 and 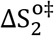 are modest (7 kcal, 9 eu). These values include contributions from the thermodynamics of UTP binding, including stacking of UTP on ATP, and from the intrinsic activation quantities for the catalytic step, but do not include any contributions from a translocation step.

Overall activation enthalpies and entropies of the next two steps (steps 3 and 4 of Mechanism 2) which begin with translocation, including opening one downstream base pair, are similar to one another and larger than those of pppApU synthesis by ~ 15 kcal and ~ 45 eu, respectively. We propose that these activation enthalpy and entropy differences arise from the translocation step. A substantial part of the positive ~ 15 kcal enthalpy of translocation is presumably opening and unstacking one downstream bp, with an enthalpy cost of 5-15 kcal/mol in solution depending on the extent of unstacking the bases in the open strands (39). The large positive activation entropy (~ 45 eu), most of which also appears to originate from the translocation step, is not simply explained because the entropy increase from base pair disruption in the initiation complex is presumably not as large as that of base pair disruption in solution (~ 25 eu).

By comparison the next five steps, all of which also begin with translocation, exhibit significantly smaller positive activation enthalpies than steps 3 and 4. The first five of these steps also exhibit negative activation entropies (see Table S3). These include the steps previously identified as ones in which contacts of the discriminator and −10 strands with RNAP are disrupted, freeing the initiation bubble strands. Most notably, the activation enthalpy and entropy of step 9 (Mechanism 2; ~ −9 kcal and −52 eu) are 30 kcal and 100 eu less than those of steps 3 and 4. Activation enthalpies and entropies for steps 5 to 8 are ~ 20 kcal and ~ 60 - 70 eu less than those of steps 3 and 4, and for step 10 are ~ 10 kcal and ~ 35 eu less than those of steps 3 and 4. The activation enthalpy and entropy for step 10 are ~10 kcal and ~35 eu less than those of steps 3 and 4, reductions which appear significant but are not as large as for the preceding steps. Four of these steps (5, 7, 8, 9) also have small rate constants k_I_. Unfavorable (negative) 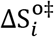 activation barriers are responsible for these small rate constants, not large positive 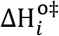.

Both the activation enthalpy and entropy of the last initiation step (step 11) are much larger than those of the preceding seven steps, though not quite as large as for steps 3 and 4. Contacts that are disrupted in this step are with the duplex (−35 and upstream) and not with single stranded DNA. Hence the unusual activation thermodynamics are confined to the steps that break RNAP contacts with the bubble strands.

### Stepwise Base Stacking of Discriminator and −10 Strands After RNAP Interactions are Disrupted

A likely source of the unusual 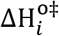 and 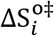 values for forming 5-mer to 10-mer RNA is base stacking interactions occurring after interactions of the strands of the discriminator and the −10 region with RNAP are disrupted in a step-wise manner by translocation stress. Enthalpies and entropies of single-strand base stacking in solution are about −5 kcal mol^−1^ and −15 eu (39); enthalpies and entropies of forming stacked base pairs in a duplex are about twice as large in magnitude. In Table S4, using these enthalpy changes, we interpret the reduced activation enthalpies of steps 5-10 as formation of 4 base stacking interactions in each of steps 5-8, 6 base stacking interactions in step 9 and 2 base stacking interactions in step 10. This totals 24 base stacking interactions, consistent with the stacking of all 11 bases of each bubble strand on their neighbors from the hybrid at +1 to the duplex at −12.

Single-stranded nucleic acid are highly stacked in solution at the temperatures investigated here. Calorimetric studies revealed that formation of the very stable wrapped SSB-ssDNA complex unstacks the bases of ssDNA, resulting in a very large positive (and temperature-dependent) binding enthalpy (40). Permanganate footprinting of the unstable RNAP-λP_R_ I_2_ open intermediate revealed that many thymines of the open strands of the initiation bubble are more stacked than in the more stable I_3_ open complex (28). Hence we expect that stepwise disruption of RNAP contacts with the discriminator and −10 strands in initiation will result in stacking of the bases in these regions.

Several aspects of the observed patterns in the rate constants and activation energies (Fig. 5; Table S3-4) are unusual. First, formation of stacked bases (as judged by small positive or negative activation enthalpies and entropies) lags the disruption of RNAP-promoter strand contacts with the discriminator and the −10 regions (as judged by small rate constants) by one step. Second, interpretation of these small activation enthalpies in terms of stacking reveals that all but the final stacking interactions within the bubble strands occur in groups of 4 or 6 bases, which is unexpected because base stacking in a single-stranded structure shows little if any cooperativity. Finally, no step has the large negative activation enthalpy and entropy that would be expected for duplex formation from these stacked strands, estimated to have a favorable enthalpy change of about −60 kcal mol^−1^. This may occur in step 9, though it would be expected in step 11 when upstream contacts are broken so there is no topological barrier to duplex formation.

The observation that no step has the large negative activation enthalpy and entropy that would be expected for duplex formation is consistent with the recent finding that reducing the stability of the bubble-region duplex by as much as 9 kcal by introducing mismatches at the promoter −6 and −10 positions has no effect on the rate of promoter escape (2). Step-by-step base stacking provides an explanation for this observation. Step-by-step stacking accompanying step-by-step disruption of RNAP-strand contacts with the discriminator and −10 regions can account for much of the favorable free energy change of duplex formation, distributing this effect over multiple steps of initiation.

## Materials and Methods

Details about reagents (buffers, enzymes, DNA), initiation kinetic assays (single-round in synthesis of full-length RNA), and analysis of amounts of transient short RNAs from productive complexes and of stalled and released short RNA from nonproductive complexes are described in reference (32) and in Supplemental.

## Acknowledgements

We gratefully acknowledge support of NIH GM R35-118100 for this research.

## Notes

### Competing Interest Statement

The authors have declared no competing interest.

### Summary of Updates

The on-figure legend of figure 5 was corrected in this revision. Point sizes in figure 5 were also adjusted for aesthetic consistency.

