## Supplemental Methods, Tables, and Figures for "Kinetic-Mechanistic Evidence for Which *E. coli* RNA Polymerase-λP_R_ Open Promoter Complex Initiates and for Stepwise Disruption of Contacts in Bubble Collapse"

### Supplemental Methods, Tables, Figures

#### Methods

##### Reagents, Buffers, and Gels

Reagents used for buffers and stock solutions were purchased in the highest available grade and used as received. All solutions were prepared using 18MΩ deionized water from a Barnstead EPure system. NTPs and dNTPs (Boston Bioproducts, Thermo Fisher, New England Biolabs) used in transcription assays and PCR reactions were 99% pure and used as received. Enzymes for PCR reactions were purchased from NEB and used according to the manufacturers protocols.

Storage Buffer for core RNAP, $\sigma^{70}$ and RNAP holoenzyme is 50% v/v glycerol, 0.01M Tris, 0.1M NaCl, 0.1 mM EDTA, and 0.1 mM DTT. Transcription buffer (TB) is 40mM Tris, 5mM $MgCl_2$ 60mM KCL, 1mM DTT, 0.05 mg/ml BSA, and is adjusted to pH 8.0 at experimental temperature (19°C, 25°C, or 37°C).

2X initiation solution (IS) for rapid quench flow (RQF) transcription assays with α-$^{32}$P-GTP is 0.1 mg/ml heparin, 400 μM ATP, 400 μM UTP, 20 μM unlabeled GTP, and 35 nM α-$^{32}$P-GTP. For assays with α-$^{32}$P-UTP IS is 0.1mg/ml heparin, 400 μM ATP, 400 μM GTP, 20 μM unlabeled UTP, and 35 nM α-$^{32}$P-UTP. The IS is mixed 1:1 with preformed OC to reach experimental concentrations of each NTP.

Quench Solution (QS) for RQF transcription assays is 8M Urea and 15 mM EDTA in TB. QS with added dyes (QSD) for polyacrylamide gel electrophorese (PAGE) gas 0.05% xylene cyanol and 0.05% bromphenol blue in QS. TBE buffer for gel electrophoresis is 90mM Tris-borate (pH 8.3) and 2mM $Na_2EDTA$. All transcription gels are 20% acrylamide-(bis)acrylamide (19:1), and were made using the UreaGel system (National Diagnostics).

##### RNA Polymerase and $\lambda P_R$ Promoter DNA

RNAP core enzyme ($\alpha_2\beta\beta'\omega$) is overexpressed, purified and assembled as previously reported (Henderson '19). Filter binding assays performed on RNAP holoenzymes used in all experiment reported here show that 50% ± 15% of RNAP molecules form a stable open complex with the $\lambda P_R$ promoter at 37°C. All RNAP concentrations reported here refer to this active fraction. $\lambda P_R$ promoter DNA is prepared as described previously (Henderson '17). Primer sequences used to assemble the $\lambda P_R$ promoter DNA fragment are reported in supplemental table S1.

##### Initiation Kinetic Assays

Initiation kinetic assays are performed and imaged as described previously (Henderson '19). These allow single-round synthesis of full-length (FL) RNA. Briefly, 2X initiation solution (IS) is mixed 1:1 with preformed OC at time zero using a KinTek Corp. Rapid Quench Flow (RQF). Reactions are quenched with QS and RNA is separated via PAGE. Both the IS and preformed OC solutions are incubated at the experimental temperature for 1 h prior to mixing, and experiments are carried out with the RQF at that same temperature (19°C, 25 °C, or 37 °C).Transcription gels are transferred to a phosphorimaging cassette and imaged via a Typhoon 9000 phosphorimager after exposure for 18 h. Results are quantified using the ImageQuant software as described previously (Henderson '17). Quantitative results reported here are from the average of 2-4 RQF experiments, and the uncertainties reported are one standard deviation from the mean.

##### Analysis of Short RNAs Synthesized by Nonproductive Complexes

The amount of FL RNA (<10-mer) and of the eight short RNA species (3-10-mer) are calculated as a function of time from phosphorimager line scans of transcription gels. The 2-mer pppApU is not labelled in $\alpha$-$^{32}$P-GTP labeled experiments and does not separate sufficiently from the unincorporated monomer band to quantify in α-$^{32}$P-UTP labelled experiments. The amount of each short RNA synthesized by nonproductive (stalled) complexes as a function of time during the fast phase of FL RNA synthesis are determined by using Origin 2018b to fit production of the first abortive RNA to a single exponential decay to a plateau as described in Henderson '19 and subtracting the predicted abortive RNA amount from the total RNA at each length as a function of time (See figure S3-S6).

**Analysis of Transient Short RNAs Synthesized by Productive Complexes**

The time dependent appearance and decay of short transient RNAs are globally fit to two mechanisms (Figure 4). Both mechanisms include reversible ATP (position 1) binding, reversible UTP (position 2), and irreversible 2-mer catalysis. All subsequent steps are described by a composite rate constant $k_i$ which describes reversible translocation, reversible NTP binding, and irreversible catalysis in one step. The two mechanisms differ in the pre-NTP binding phase. Step 1 of mechanism 1 is ATP binding. In mechanism 2 step 1 is a reversible conversion from OC to the initiation complex, denoted IC. Fitting is done in Kintek Explorer v6.3.180116 (Kenneth Johnson, Kintek Corp). The resulting rate constants are used to predict the productive short RNA amounts as a function of time to compare to experimental data.

**Commentary on Figures S3-6:**

Amounts of each detectable short RNA species (3-mer to 10-mer) formed prior to promoter escape at the low-UTP and high-UTP conditions at 25 $^{o}$C and 37 $^{o}$C are plotted vs. time (linear scale) in Figs. S3-S6. At 25 $^{o}$C, high UTP, significant transient peaks in amounts of all short RNAs except 7-mer are observed. This is consistent with previously-reported observations at 25 $^{o}$C, high UTP, high UTP (Henderson '19). Transient buildup of N-mer RNA is expected when the synthesis of N-mer is relatively rapid and the subsequent synthesis of (N+1)-mer is slow. At high UTP (and low GTP) the first step of initiation (pppApU synthesis) is rapid and not a bottleneck, and intermediates (5-mer, 9-mer, 10-mer) preceding steps where G is added are expected to accumulate because of the low GTP concentration. Accumulation of 3-mer, 4-mer, 6-mer and 8-mer must however indicate that the rate constants of the following steps of RNA synthesis (synthesis of 4-mer, 5-mer, 7-mer and 9-mer) must be small. This was previously observed at this NTP condition at 19 $^{o}$C, where rate constants for synthesis of these four RNAs as well as 8-mer and 11-mer are smaller than for other steps of initiation.

By comparison, at 25 $^{o}$C, low UTP, where synthesis of pppApU is slowed by the low UTP concentration, significant transient amounts of only three RNAs (3-mer, 6-mer, 10-mer) are observed. This result is also consistent with the previous finding at 19 $^{o}$C (Henderson '19). Analysis of the kinetics of accumulation and decay of these transient RNAs at the different NTP conditions investigated yields overall second order rate constants $k_i$ for the individual steps of NTP incorporation into the RNA-DNA hybrid at 25 $^{o}$C to compare with the published values at 19 $^{o}$C (see Discussion).

In contrast, at the high UTP condition at 37°C only four transient RNA intermediates (5-mer, 6-mer, 9-mer, 10-mer) from productively-initiating complexes are detected (Fig. S6), and no transient RNAs are detected at the low UTP condition (Fig. S5). These observations are consistent with the proposal above that an additional step at the beginning of the mechanism slows the kinetics of FL RNA synthesis at 37 °C, especially at the low UTP condition.

**Determination of Activation Thermodynamics for Individual Steps of Initiation**

Values of ln $k_i$ for each step of initiation are plotted vs. 1/T in Fig. S7. Arrhenius activation energies $E_A$ calculated from the slopes (slope = $-E_A/R$) are plotted in Fig. 5 and listed in Table S3. Activation free energies ($\Delta G_i^{o\ddagger}$) for each step of RNA-DNA hybrid extension are calculated from the equation

$$\ln k_i = \ln k_{max} - (\Delta G_i^{o\ddagger}/RT) \qquad \text{Eq. S1}$$

where $\Delta G_i^{o\ddagger}$ is the free energy difference between the high free energy transition state of the catalytic step and the reactants (incoming NTP, pre-translocated state of the initiation complex) and $k_{max}$ is the maximum rate constant for the hypothetical situation where $\Delta G_i^{o\ddagger} = 0$. Because $k_{max}$ is a second order rate constant, we estimate it as a diffusion-limited rate constant for NTP binding ($k_{max} \approx 10^{10}$ $M^{-1}$ $s^{-1}$). The uncertainty in this estimate, though large, is inconsequential for the analysis reported here. We assume $k_{max}$ is the same for each step of initiation and treat it as temperature-independent. If $k_{max}$ is a diffusion-limited rate constant, its temperature dependence is not sufficiently large to be significant in the analysis reported here.

From $k_i$ values as a function of temperature (Fig. S7) and $k_{max}$, activation free energies and their enthalpic and entropic components are obtained for each initiation step:

$$\Delta G_i^{o\ddagger} = -RT \ln (k_i/k_{max}) = \Delta H_i^{o\ddagger} - T\Delta S_i^{o\ddagger}$$

where
$$\Delta H_i^{o\ddagger} \approx E_A = -R\left(\frac{\partial \ln k_i}{\partial\left(\frac{1}{T}\right)}\right) \text{ and } \Delta S_i^{o\ddagger} = (\Delta H_i^{o\ddagger} - \Delta G_i^{o\ddagger})/T \qquad \text{Eq. S2}$$

Values of $\Delta G_i^{o\ddagger}$, $\Delta H_i^{o\ddagger}$, $\Delta S_i^{o\ddagger}$ and their uncertainties for each step of extension of the RNA-DNA hybrid by NTP incorporation at 19 °C are listed in Table S3.

**Table S1: Primers used for λ$P_R$ template preparation**

| | |
|---|---|
| λ$P_R$_wt_forward (-71 to -12) | CCACGAATTCGGATAAATATCTAACACCGTGCGTGTTGACTATTTTACCTCTGGCGGTG |
| λ$P_R$_wt_reverse (-24 to 31) | ACAAAACCTTCATAGAACCTCCTTACTACATGCAACCATTATCACCGCCAGAGGT |
| HBOT | CACCTGCACCGACAAAACCTT |
| HTOP | CCAGCATTCCTCCACGAATTC |

**Table S2: Composite Rate Constants $k_i$ ($\mu M^{-1}$ $s^{-1}$) for Each Initiation Step at 19, 25, and 37 °C**

| Base Position i | Base Incorporated | 19 °C | 25 °C | 37 °C [a] |
|---|---|---|---|---|
| 2 | U | 0.63 ± 0.22 | 0.69 ± 0.24 | 1.5 ± 0.5 |
| 3 | G | 0.15 ± 0.08 | 0.49 + 0.64[b] | > 2 |
| 4 | U | 0.012 ± 0.001 | 0.033 ± 0.005 | > 0.2 |
| 5 | A | 0.064 ± 0.044 | 0.077 ± 0.040 | 0.11 ± 0.07 |
| 6 | G | 0.16 ± 0.04 | 0.21 ± 0.04 | 0.24 ± 0.08 |
| 7 | U | 0.017 ± 0.002 | 0.021 ± 0.003 | > 0.03 |
| 8 | A | 0.037 ± 0.020 | 0.044 ± 0.022 | > 0.07 |
| 9 | A | 0.052 ± 0.035 | 0.043 ± 0.021 | 0.033 ± 0.024 |
| 10 | G | 0.13 ± 0.03 | 0.22 + 0.61[b] | 0.62 ± 0.17 |
| 11 | G | 0.028 ± 0.003 | 0.049 ± 0.006 | 0.21 ± 0.03 |

[a]Uncertainties in 37 °C $k_i$ values could not be determined from the fit because of the insufficient number of detectable small RNA transient intermediates (Figs S5-6). Manual variation of parameters in Kintek Explorer showed that for steps with no detected transients (3-mer, 4-mer, 7-mer, 8-mer) these $k_i$ values are lower bounds, in that use of larger $k_i$ values for these steps did not affect the fit. When combined with well-determined 19 °C and 25 °C $k_i$ values, these 37 °C lower-bound $k_i$ values give linear Arrhenius plots (Fig. S7), indicating their validity. Uncertainties in other 37 °C $k_i$ are estimated using the larger of the uncertainties for that step in the 19 °C and 25 °C data.

[b] Relatively large uncertainties in these 25 °C $k_i$ values reflect insufficient detectable intermediates to fit in the Kintek Explorer analysis. Cited uncertainties are estimated from the range of acceptable $k_i$ values for these steps. For the 9-10 step, use of this this 25 °C $k_i$ value together with better-determined 19 °C and 37 °C $k_i$ values yields a linear Arrhenius plot (Fig. S7), indicating its validity.

**Table S3: Arrhenius Activation Energies $E_{A,i}$ and Transition State Barriers $\Delta G_i^{o\ddagger}$, $\Delta H_i^{o\ddagger}$, $\Delta S_i^{o\ddagger}$ for Each Step of Initiation**

| Base Position i | $k_I$ ($\mu M^{-1}\ s^{-1}$) | $\Delta G_i^{o\ddagger}$ (kcal/mol)[a] | $E_{A,i}$ (kcal/mol) | $\Delta H_i^{o\ddagger}$ (kcal/mol)[a] | $\Delta S_i^{o\ddagger}$ (eu)[a] |
|---|---|---|---|---|---|
| 2 | 0.63 ± 0.22 | 4.3 ± 0.3 | 9 ± 5 | ~ 7 | ~ 9 |
| 3 | 0.15 ± 0.08 | 5.1 ± 0.8 | 27 ± 14 | ~ 22 | ~ 58 |
| 4 | 0.012 ± 0.001 | 6.6 ± 0.1 | 26 ± 2 | ~ 21 | ~ 49 |
| 5 | 0.064 ± 0.044 | 5.6 ± 0.3 | 5.1 ± 10 | ~ 0 | ~ -19 |
| 6 | 0.16 ± 0.04 | 5.1 ± 0.1 | 7.2 ± 3 | ~ 2 | ~ -10 |
| 7 | 0.017 ± 0.002 | 6.4 ± 0.1 | 6.8 ± 2 | ~ 2 | ~ -16 |
| 8 | 0.037 ± 0.020 | 5.9 ± 0.3 | 6.6 ± 7 | ~ 2 | ~ -15 |
| 9 | 0.052 ± 0.039 | 5.7 ± 0.3 | -4.4 ± 10 | ~ -9 | ~ -52 |
| 10 | 0.13 ± 0.03 | 5.2 ± 0.2 | 16 ± 4 | ~ 11 | ~ 20 |
| 11 | 0.028 ± 0.003 | 6.1 ± 0.1 | 21 ± 2 | ~ 16 | ~ 34 |

[a] $\Delta G^{o\ddagger}, \Delta H^{o\ddagger}, and\ \Delta S^{o\ddagger}$ are defined in Eq. S1-S2.

**Table S4: Interpretation of Rate Constants $k_i$ and Barriers $\Delta G_i^{o\ddagger}$, $\Delta H_i^{o\ddagger}$, $\Delta S_i^{o\ddagger}$ for Each Step of Initiation**

| Base Position i | $k_i/k_3$ | Region, Fraction Of Contacts Disrupted [a] | Proposed Stacked Bases in Bubble [b] |
|---|---|---|---|
| 2 | ~ 4.2 | - | 0 |
| 3 | 1 | - | 0 |
| 4 | ~ 0.1 | Disc; ¾ | 0 |
| 5 | ~ 0.4 | Disc; ¼ | Disc; 4 |
| 6 | ~ 1 | - | Disc; 4 |
| 7 | ~ 0.1 | -10; ½ | Disc; 4 |
| 8 | ~ 0.2 | -10; ¼ | -10; 4 |
| 9 | ~ 0.3 | -10; ¼ | -10; 6 |
| 10 | ~ 0.9 | - | -10; 2 |
| 11 | ~ 0.2 | -35; all | 0 |

[a] Fractions of contacts in a region disrupted in step i, estimated from ($\Delta G_i^{o\ddagger}$- $\Delta G_3^{o\ddagger}$)/Σ($\Delta G_i^{o\ddagger}$- $\Delta G_3^{o\ddagger}$) where i = 4, 5 for discriminator (abbreviated Disc) region and i = 7, 8, 9 for -10 region. Similar results are obtained using $\Delta G_4^{o\ddagger}$ instead of $\Delta G_3^{o\ddagger}$ as the reference.

[b] For steps 5 – 9, the number of stacked bases is estimated from ($\Delta H_3^{o\ddagger}$- $\Delta H_i^{o\ddagger}$)/5, assuming -5 kcal per mol per base stacking interaction. Similar results are obtained from analysis of ($\Delta S_3^{o\ddagger}$- $\Delta S_i^{o\ddagger}$) assuming an entropy change of approximately – (15 – 20) eu per stacking interaction. For both enthalpy and entropy analyses, similar results are obtained using $\Delta H_4^{o\ddagger}$ instead of $\Delta H_3^{o\ddagger}$ and $\Delta S_4^{o\ddagger}$ instead of $\Delta S_3^{o\ddagger}$ as the reference.

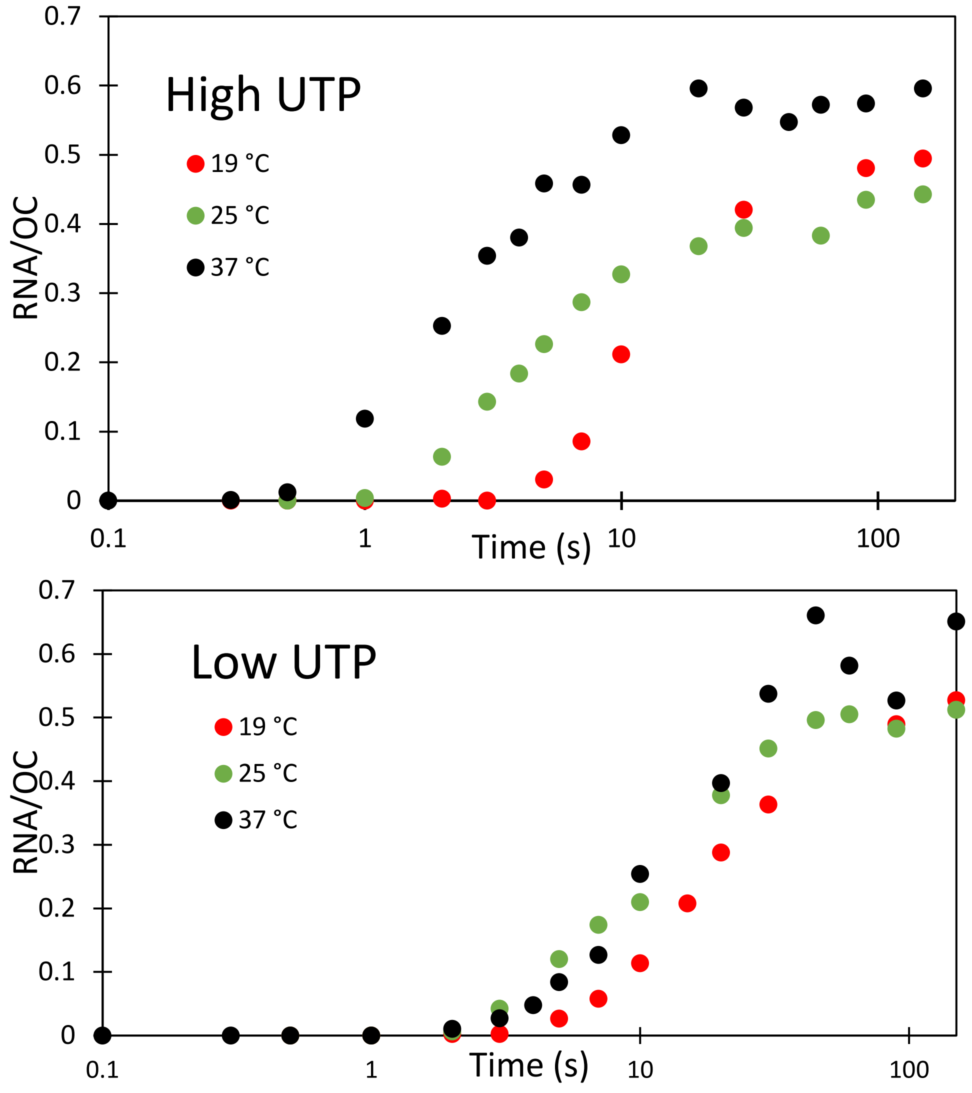


**Figure S1: Kinetics of FL RNA Synthesis in Single-Round Initiation from LPR Promoter as a Function of Temperature.** Amounts of FL RNA (>10 bp) synthesized as a function of time at 19 °C (red), 25 °C (green), and 37 °C (black), normalized per open complex (RNA/OC). Data are from individual representative experiments at the low GTP condition (10 µM GTP, 200 µM ATP, UTP (top)) and the low UTP condition (10 µM UTP, 200 µM ATP, GTP (bottom)). Experiments are performed at a 2:1 ratio of active RNAP to promoter DNA, where 100% of promoter DNA forms a stable OC at these temperatures.

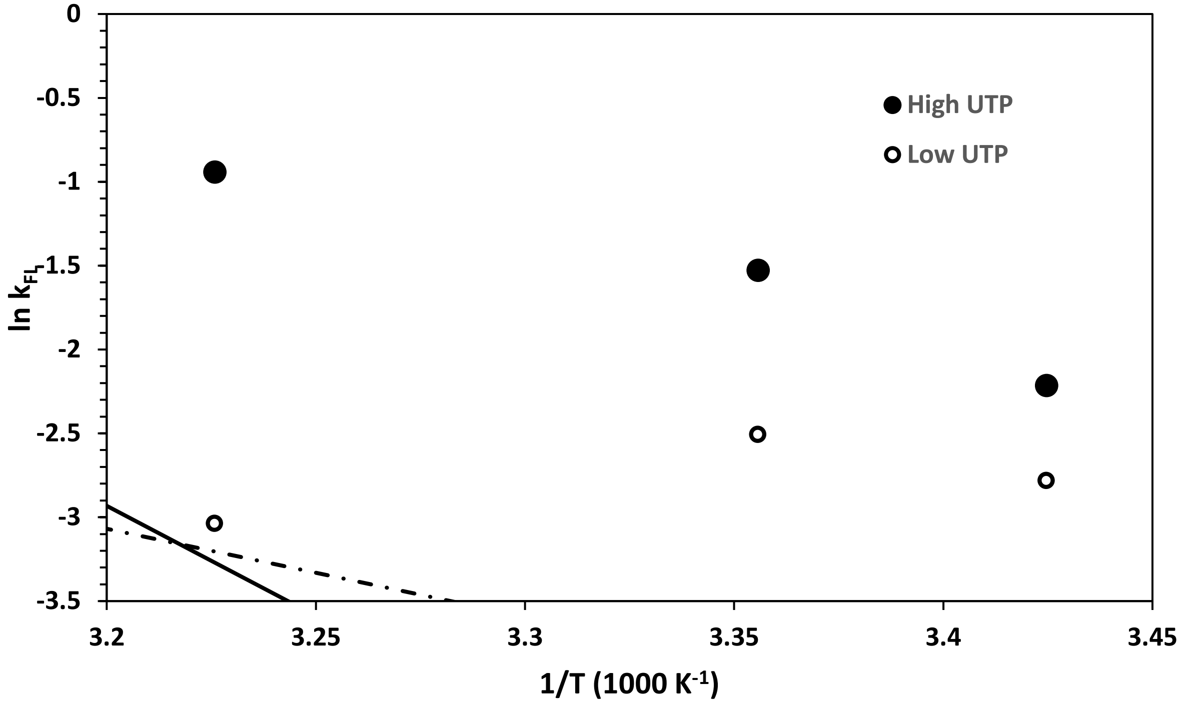


**Figure S2: Arrhenius Plot of ln$k_{FL}$ vs 1/T.** Plot of the natural logarithm of the first order decay rate constant for full length RNA production ($k_{FL}$, in s$^{-1}$) vs inverse temperature (K$^{-1}$) for both 10 µM GTP (High UTP, filled circles) and 10 µM UTP (Low UTP, open circles). For both conditions, and particularly low UTP, the 37 °C rate constant is significantly smaller than predicted assuming an Arrhenius relationship between the 19 °C and 25 °C rate constants (predictions shown).

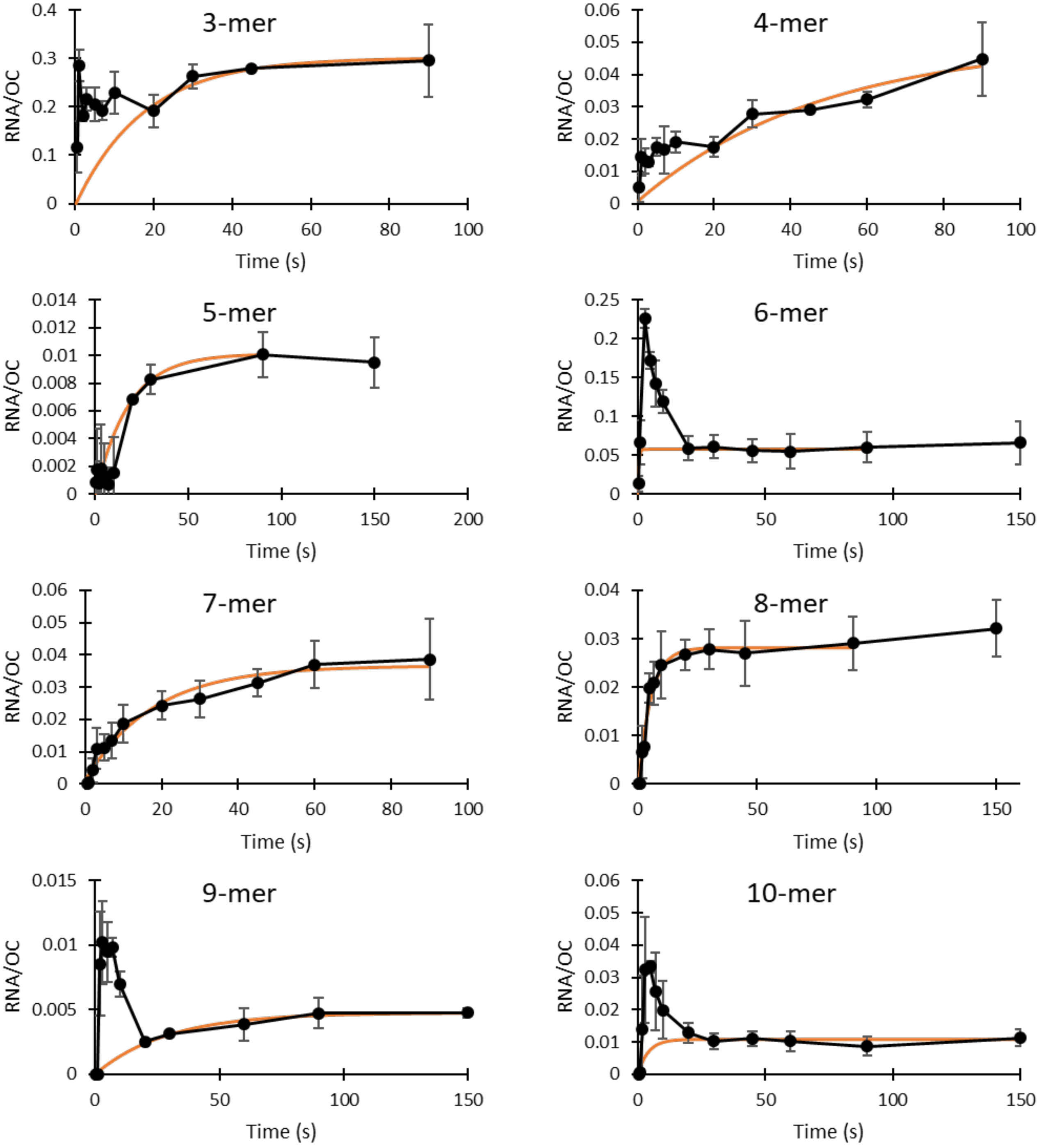


**Figure S3: Average Amounts of Short RNAs vs Time at 25 °C, Low UTP.** Amounts of 3-mer to 10-mer RNA per OC (averages of 2-4 independent experiments) are shown with one standard deviation error bars (black). Data for all detectable lengths up to the point of escape are shown. NTP concentrations are 200 µM ATP and GTP, 10 µM UTP. For each RNA length, the predicted contribution of the non-productive fraction of OCs is also shown (orange).

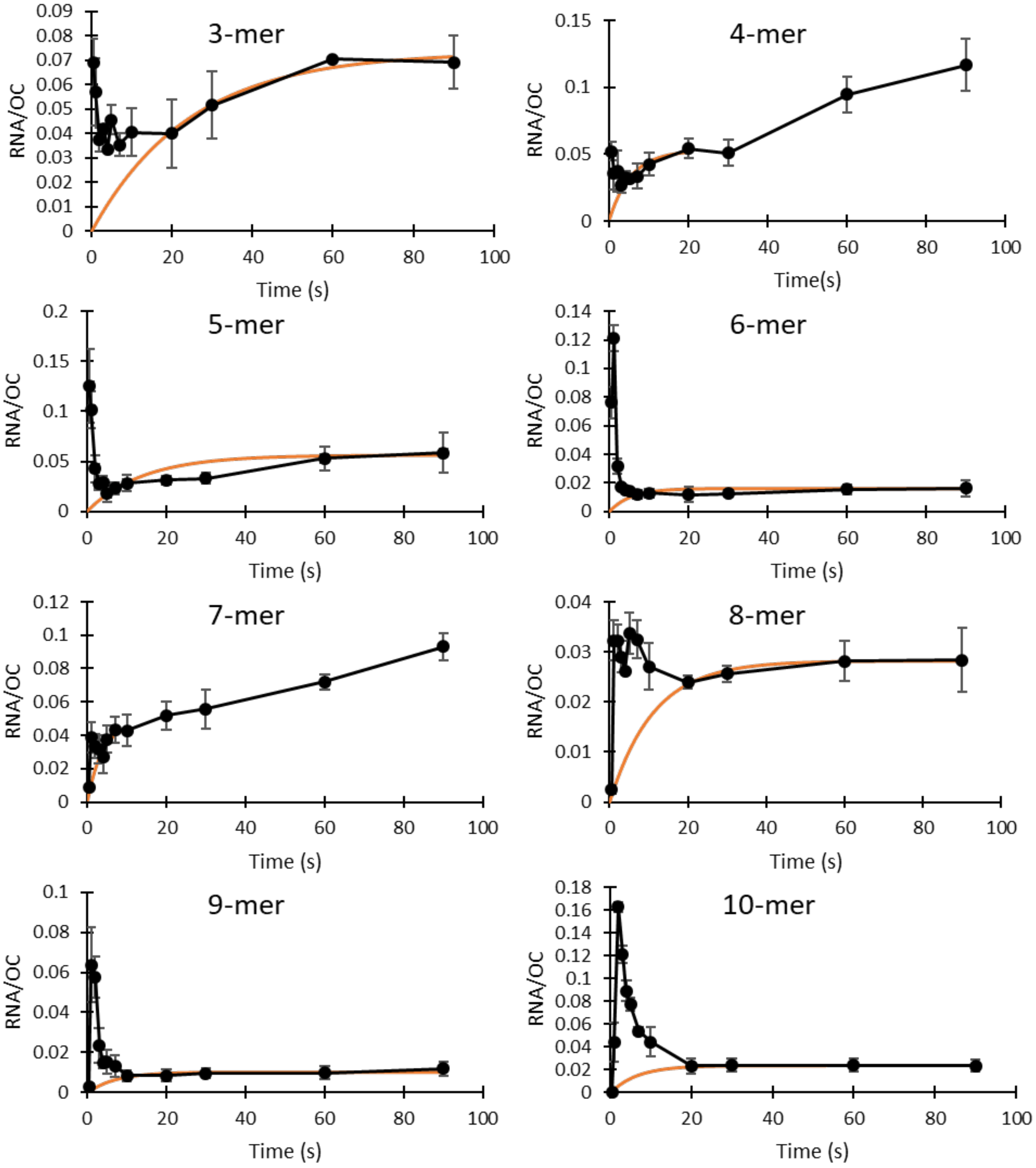


**Figure S4: Average Amounts of Short RNAs vs Time at 25 °C, High UTP.** Amounts of 3-mer to 10-mer RNA per OC (averages of 2-4 independent experiments) are shown with one standard deviation error bars (black). Data for all detectable lengths up to the point of escape are shown. NTP concentrations are 200 µM ATP and UTP, 10 µM GTP. For each RNA length, the predicted contribution of the non-productive fraction of OCs is also shown (orange).

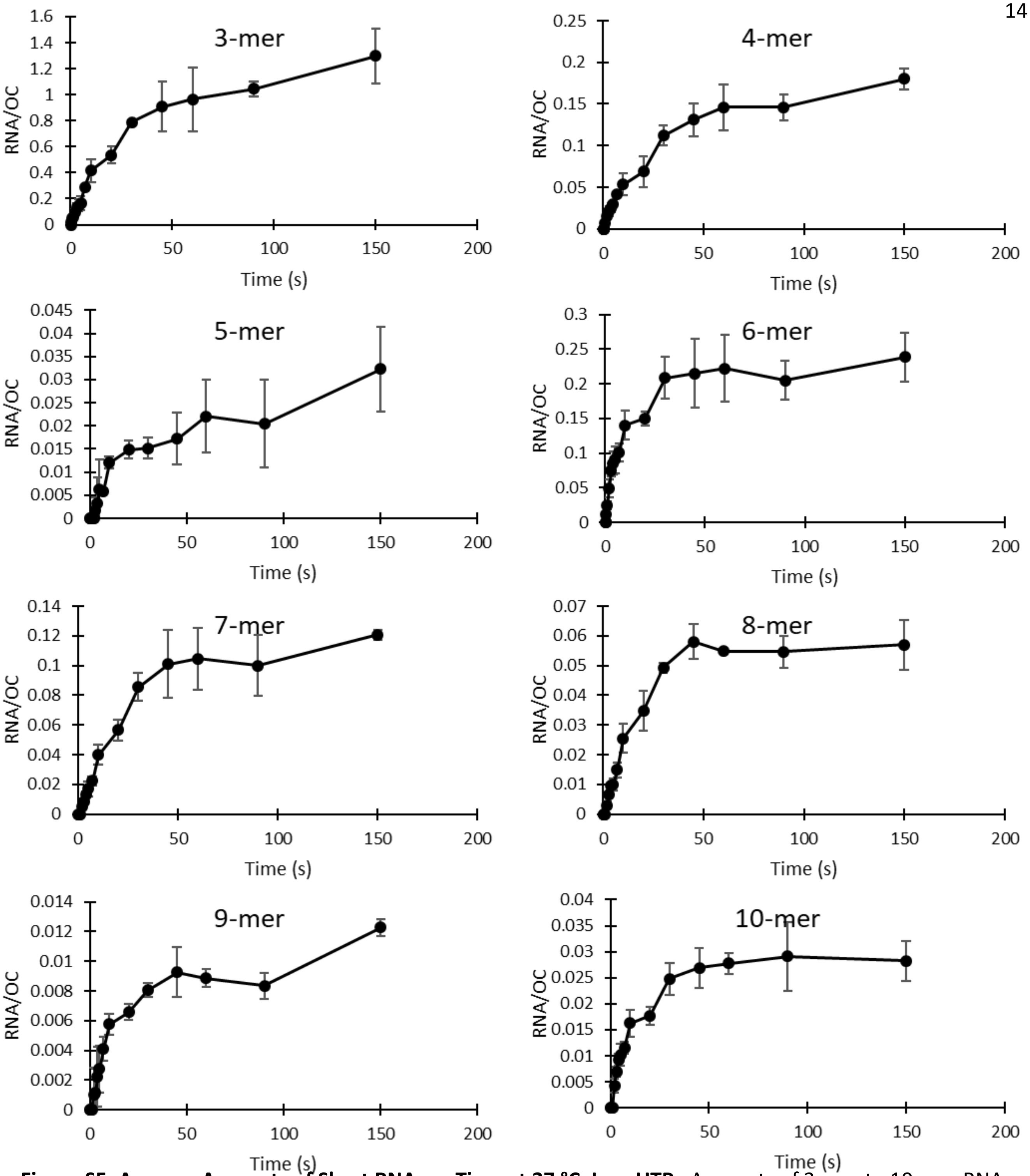


**Figure S5: Average Amounts of Short RNAs vs Time at 37 °C, Low UTP.** Amounts of 3-mer to 10-mer RNA per OC (averages of 2-4 independent experiments) are shown with one standard deviation error bars (black). Data for all detectable lengths up to the point of escape are shown. NTP concentrations are 200 µM ATP and GTP, 10 µM UTP. Transient concentrations of short RNA synthesized by productive OC are apparently too small to detect at 37 °C, low UTP, and therefore these curves show the initial and subsequent rounds of RNA synthesis by the non-productive fraction of OCs.

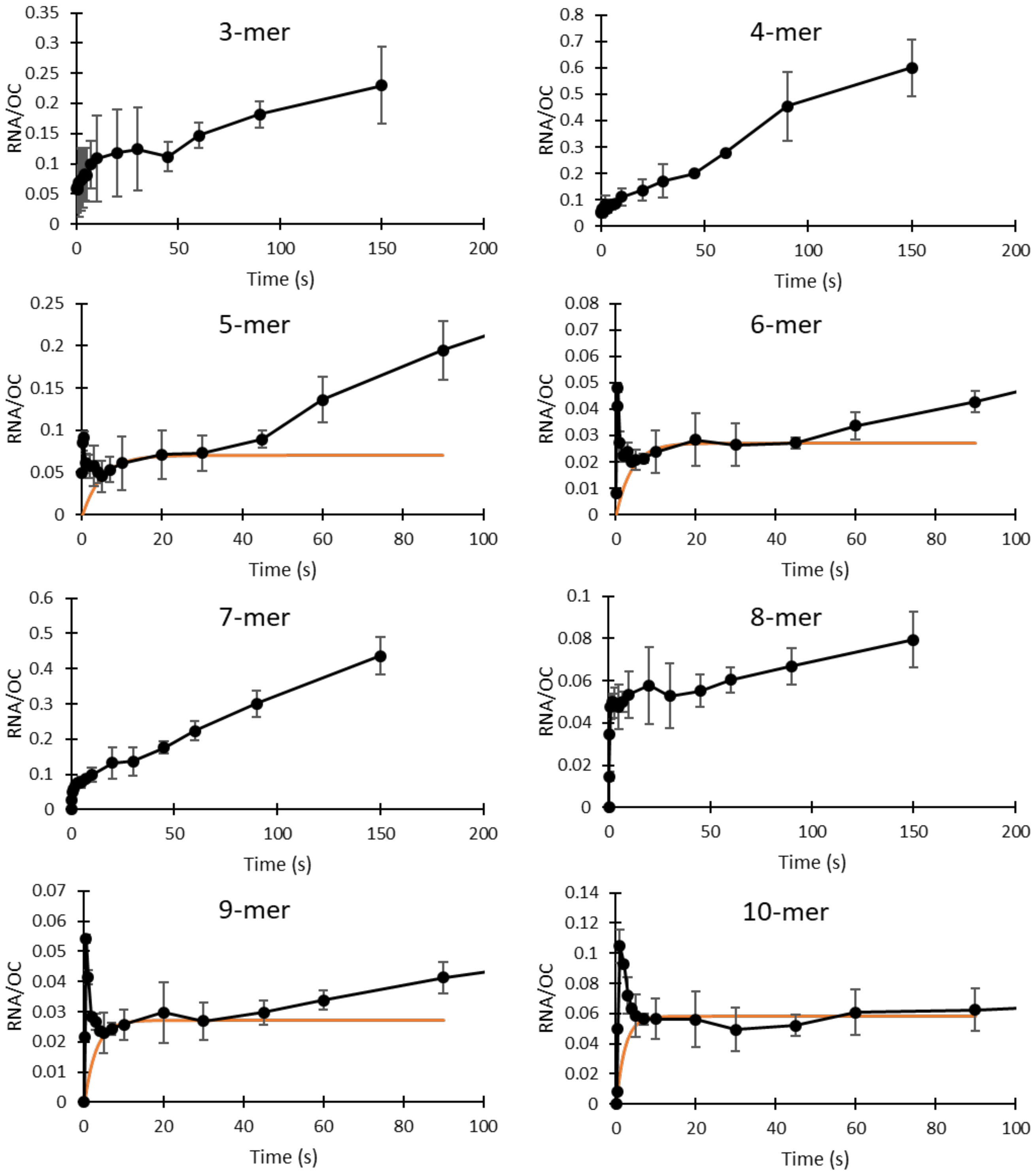


**Figure S6: Average Amounts of Short RNAs vs Time at 37 °C, High UTP.** Amounts of 3-mer to 10-mer RNA per OC (averages of 2-4 independent experiments) are shown with one standard deviation error bars (black). Data for all detectable lengths up to the point of escape are shown. NTP concentrations are 200 μM ATP and UTP, 10 μM GTP. For each RNA length where a transient concentration of RNA from synthesis by productive OC can be detected, the predicted contribution of the non-productive fraction of OCs is also shown (orange).

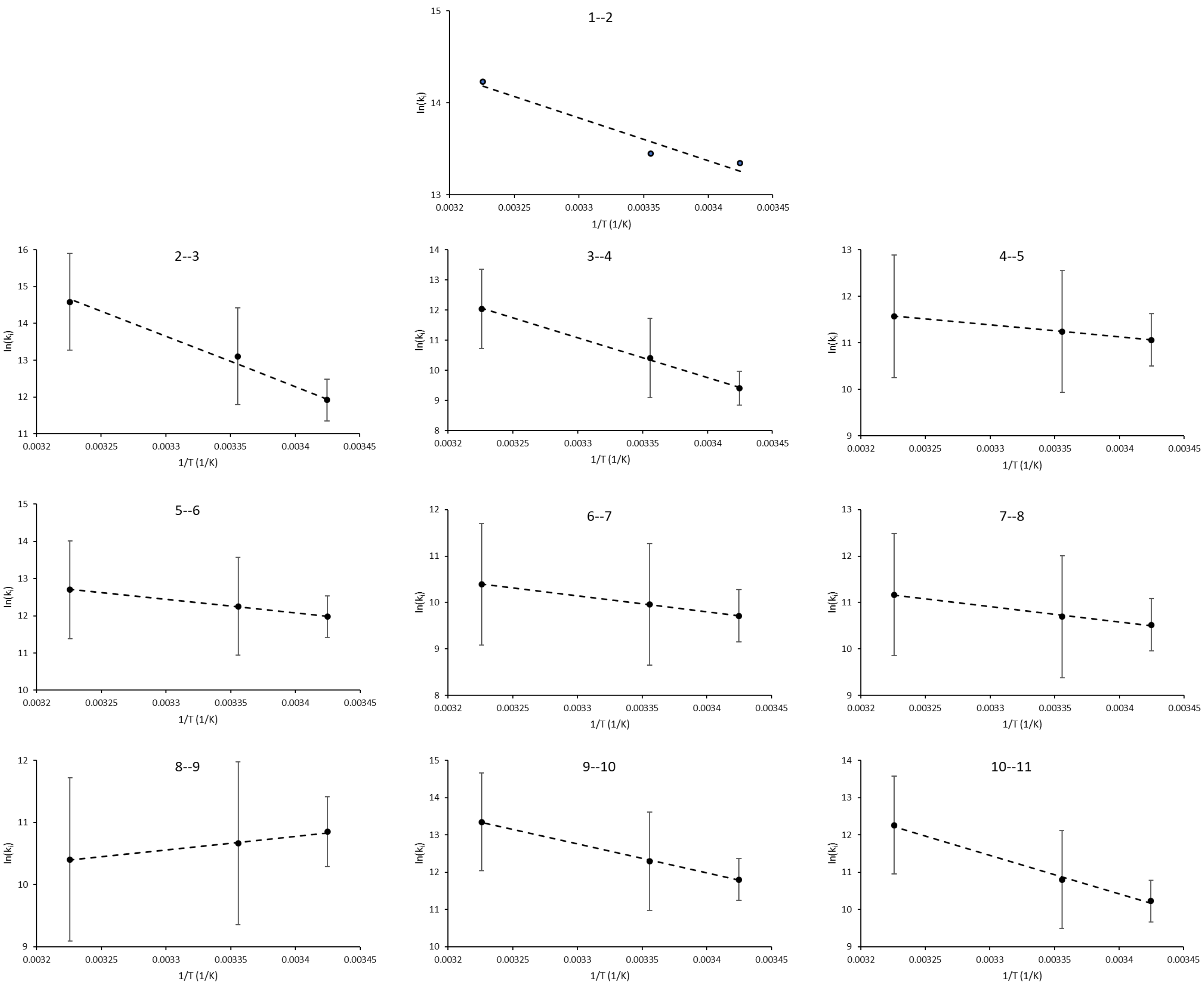


**Figure S7: Temperature Dependence of Composite Initiation Rate Constants $k_i$: Arrhenius Plots**

For each step, values of $\ln k_i$ (with $k_i$ in $M^{-1}s^{-1}$) are plotted vs 1/T. Uncertainties are listed in Table S2. Slopes of linear fits yield Arrhenius activation energies $E_A$ (slope = $-E_A/R$), reported in Table S3.
